# Tiny killers: first record of rhabdocoel flatworms feeding on water flea embryos

**DOI:** 10.1101/2025.02.10.636452

**Authors:** Nedim Tüzün, Nina Lemke, Yander L. Diez, Tom Artois, Marlies Monnens

## Abstract

Flatworms are increasingly recognized for their ecological significance and potential to disrupt local fauna, yet most research has focused on conspicuous, larger planarians. Smaller flatworms, or microturbellarians, are often top predators within meiofaunal food webs. Here, we report a novel interaction involving a rhabdocoel microturbellarian, *Strongylostoma simplex*, predating on *Daphnia* water flea embryos. We identified the flatworm based on histological serial sections and recognized key diagnostic traits. In a laboratory experiment, we tested for survival and offspring production of *Daphnia magna* in the presence and absence of *Strongylostoma simplex*. Exposure to flatworms caused a drastic reduction in water flea fitness, indicated by the strongly reduced survival and offspring production in flatworm-exposed *Daphnia*. This finding corroborates our visual observations of egg predation by these flatworms, and suggests a strong pressure on *Daphnia* population dynamics. This is particularly concerning for small or isolated water bodies, such as the water reservoirs located in a cemetery in Berlin in which we documented this interaction, as this would increase the probability of encounters between flatworms and water fleas. As *Daphnia* play an essential role in regulating phytoplankton blooms and supporting higher trophic levels in freshwater ecosystems, understanding the ecological consequences of potentially invasive predatory flatworms is imperative.

## INTRODUCTION

In recent years, a growing body of research has documented the rise of invasive flatworms, particularly large land planarians that have been increasingly reported across Europe (Álvarez-Presas et al., 2014; Breugelmans et al., 2012; de Waart, 2016; Jones et al., 2019; Justine et al., 2014, 2020; Lago-Barcia et al., 2019; Mazza et al., 2016; Sluys, 2016; Soors et al., 2019; Vardinoyannis & Alexandrakis, 2019). It is becoming increasingly evident that these flatworms exert large ecological impacts, particularly through their carnivorous feeding habits that target local fauna (Boag and Neilson, 2006; de Waart, 2016; Sluys, 2016). Moreover, they may indirectly affect ecosystem functioning by influencing soil fertility and drainage (Sluys, 2016). While the invasions and ecological processes involving large land planarians are becoming better understood, the role of smaller flatworms, collectively referred to as microturbellarians, remains largely understudied. Such meiofaunal organisms are, however, known to play key roles in several ecosystem processes and services (Bonaglia et al., 2014, 2020; Piot et al., 2014; Schratzberger & Ingels, 2018).

A prime example of this is Rhabdocoela, the most species-rich taxon of microturbellarians (WoRMS, 2024). As top predators within meiofaunal food webs, these animals likely play critical roles in such ecosystems. Although a substantial body of literature exists on rhabdocoel ecology, predation behaviour, and dietary preferences, much of this research dates back several decades and focuses primarily on few mesostomid species (Blaustein & Dumont, 1990; Brendonck et al., 2002; Case & Washino, 1979; De Roeck et al., 2005; De Meester & Dumont, 1990; Dumont & Carels, 1987; Dumont et al., 2014; Dumont & Schorreels, 1990; Jennings, 1957; Kaur, 2000; Kolasa, 1984; Kolasa & Schwartz, 1988; Maly et al., 1981; Menn & Armonies, 1999; Rocha et al., 1990; Schwartz & Hebert, 1982, 1986; Tranchida et al., 2009; Wrona & Koopowitz, 1998). However, several of these studies already indicate that rhabdocoel flatworms can alter invertebrate community structures through predation pressure (Blaustein, 1990; Blaustein & Dumont, 1990; Case & Washino, 1979; Maly et al., 1981; Schwartz & Hebert, 1982; Tranchida et al., 2009).

For those rhabdocoels whose diet is known, cladoceran (Crustacea) zooplankton appear to be a common prey (Blaustein & Dumont 1990; Dumont et al., 2014; Kolasa & Schwartz, 1988; Rocha et al., 1990). Large zooplankton such as the water flea *Daphnia* are key ecological interactors in freshwater food webs, as they efficiently graze on phytoplankton and are preferred prey for a range of predators (Miner et al., 2012). Studies reveal that flatworms can exert strong pressure on cladocerans, with effects ranging from shortened lifespan (Nandini & Sarma 2013) to reduced population size (Caramujo & Boavida 2000; Maly et al., 1981; Wang et al., 2011), and ultimately altered community structure and ecosystem functioning (Devkota et al., 2023). Research efforts on flatworm predation on cladocerans has almost explicitly focused on species of *Mesostoma* (Blaustein & Dumont 1990; Dumont et al., 2014; Rocha et al., 1990), and only a handful of studies on the interactions between non-mesostomid flatworms and cladocerans exists (see Houben et al., 2014; Nandini & Sarma, 2013, Tessens et al., 2023; Wang et al., 2011). Flatworms can rapidly reach high population densities in small water bodies (Blaustein & Dumont, 1990), where cladocerans such as *Daphnia* typically occur. Therefore, the investigation of novel predator-prey interactions is critical for understanding the broader ecological impacts of flatworms on freshwater community dynamics.

In this study, we report a potential new invasion of microturbellarian flatworms in Berlin, Germany. The location of this invasion has been regularly sampled for water fleas, and we recently observed a sudden, dense population of rhabdocoel flatworms, both inside brood chambers of water fleas as well as free-swimming (AFL, NL & NT, personal observation), previously unrecorded in the area. The species is identified through morphological study, and its taxonomic status is re-assessed. In addition, the potential impact of this invasion on local *Daphnia* water flea populations is explored through a short-term *in vitro* experiment.

## MATERIAL & METHODS

### Flatworm specimens and sampling location

Flatworm specimens used in the morphological study were collected from a single water well in a cemetery in Berlin (52°30’59.1”N, 13°16’56.9”E) during September 2023. We measured basic environmental parameters of the well water (temperature, pH, conductivity, dissolved oxygen) during September 2024, using a WTW Multi3630 probe.

### Morphological study

Specimens of *Strongylostoma simplex* selected for morphological study were transported to the Diepenbeek campus of Hasselt University, where they were fixed in hot Bouin’s fixative at 50°C and embedded in paraffin. The samples were then serially sectioned at 4 µm using a Leica SM2000 R Microtome in sagittal, frontal, and horizontal planes. The sections were stained with Heidenhain’s haematoxylin and counterstained with erythrosine.

A Leica LED DM2500 microscope, equipped with a drawing mirror, was used to study the sections and create a reconstruction of the internal organs. Micrographs and measurements were taken using the LAS X software provided by the supplier, with measurements performed along the central axis of the studied structures. To the authors’ knowledge, no type material for this species, nor either of its two subspecies, exists for comparative study.

### Observations of the flatworm – water flea interaction

Following the accidental observation of flatworm predation in *Daphnia magna* water flea specimens collected from the cemetery well, we first conducted *in situ* field observations. To understand the behavior of *Strongylostoma simplex* in the presence of *Daphnia magna*, and vice versa, we made live observations by adding specimens of *Daphnia* and flatworms into transparent containers filled with water. Water fleas were collected from the same cemetery well, as well as from additional cemeteries in Berlin in which we detected flatworm-infected water fleas. For a more detailed observation, we placed water fleas and flatworms in a Petri dish under a camera-stereomicroscope set-up (Olympus DP23 camera mounted on an Olympus SZX16 stereomicroscope). We observed infected water fleas (i.e. with flatworms present in the brood chamber), as well as uninfected water fleas in the presence of (free-swimming) flatworms. Observations under the stereomicroscope were recorded in photo and video format.

### Experimental design to test effects of flatworms on water fleas

To test for the effects of the flatworms on water fleas, we performed a short-term experiment where we exposed individual water fleas to flatworms, and measured two fitness-related traits: water flea survival and offspring production. We used 8 replicates per the two treatments, i.e. control (no flatworm) and flatworm-treatment (total N=16). Each water flea was housed individually in 300 ml vials filled with tap water, and fed every third day with dry yeast. Vials were refreshed every third day. We checked for survival and hatched offspring every second day. We recorded survival and total number of offspring produced over the 9-day experimental period.

For the treatment group, we added one individual water flea per vial that contained flatworms in its brood chamber. We visually confirmed the presence of flatworms in the water flea brood chamber, but did not count the number of flatworms per water flea. For the control group, we added one individual water flea per vial that did not contain any flatworms in its brood chamber. Individual water fleas were selected to be similar in size, and all carried eggs at the beginning of the trial (eggs were of similar developmental stage). During the experiment, conducted in early October, vials were exposed to the natural day-night regime (ca. 12:12 light:dark) and standard room temperature (between ca. 19–22°C).

### Statistical analyses of experimental data

To test for differences in water flea survival between flatworm-treated and the control group, we used Fisher’s exact test. To test for differences in offspring production between flatworm-treated and the control group, we used the Wilcoxon rank-sum test. This non-parametric test is preferred for data that deviate from assumptions of normal distribution and homogeneity of variances. All analyses were performed in R version 4.3.2 (R Core Team, 2023).

**Abbreviations used in figures:** bg: basophilic gland; br: brain; c: cilia; cga: common genital atrium; de: ejaculatory duct; eg1: eosinophilic gland 1; eg2: eosinophilic gland 2; epg: epidermal gland; gp: gonopore; od: oviduct; ov: ovary; pc: prepharyngeal cavity; ph: pharynx; rs: seminal receptacle; t: testis; vd: vitelloduct; vdf: vas deferens; vi: vitellaria; vs: seminal vesicle

## RESULTS & DISCUSSION

### Taxonomical account

**Dalytyphloplanida Willems, Wallberg, Jondelius, Littlewood, Backeljau, Schockaert & Artois, 2006**

**Neotyphloplanida Willems, Wallberg, Jondelius, Littlewood, Backeljau, Schockaert & Artois, 2006**

**Limnotyphloplanida Van Steenkiste, Tessens, Willems, Backeljau, Jondelius & Artois, 2013**

**Typhloplanidae Graff, 1905**

***Strongylostoma simplex simplex* Meixner, 1915**

**Figs. 1–3**

#### New locality

Luisenkirchhof II in Berlin, Germany (52°30’59”N, 13°16’57”E). Water reservoir made out of concrete (diameter 75 cm, height 70 cm) in a cemetery, with ca. 1 cm sediment on the bottom (Appendix Fig. S1), filled with tap water (no natural water flow), and frequently used as a source of drinking water by wild animals. Habitat includes water fleas (*Daphnia magna* and *D. longispina*), diving beetles, larvae of mosquitoes and mayflies. Filamentous algae were present. Water parameters: 23.0 °C, 7.823 pH, 677 µS/cm conductivity, 7.60 mg/l dissolved oxygen (measured on 7 September 2024).

#### Previously known distribution

Lunzer See (Meixner, 1915) and Schwarzensee, Austria (Steinböck, 1926), Lago Maggiore, Italy (Steinböck, 1948, 1949, 1951), Lake Mývatn, Iceland (Steinböck, 1948), and Baraus Lake, Tsjeljabinsk, Russia (Rogozin, 2017). Luther (1963) also references a record by Steinböck (1932) of the species occurring in Lago di Como and Lago di Garda, Italy, which we were unable to confirm. Note that, according to Luther (1963), it is uncertain whether the historical records prior to his work in 1963 pertain to *Strongylostoma simplex simplex* or *Strongylostoma simplex lapponicum* Papi in Luther, 1963.

#### Material

Video recordings and photographs of live specimens. Six serial sections: three in sagittal orientation, one in frontal, and two in transverse orientation.

#### Description

The specimens are 0.31–0.51 mm long (n=2), with a width approximately half the length of the body. Both the anterior and posterior ends of the body are smoothly rounded. Two brown-pigmented eyes are located at the anterior end. The epidermis is cellular and fully ciliated (Figs. 1E–F: c) and measures approximately 10 μm in height (n=2). The cilia measure approximately 7 μm in two specimens. Circular and longitudinal muscles are present above the basal lamina. The animal is predominantly brown, except for its transparent edges and a light-colored anterior region. The eggs of the animals, approximately 200 μm in diameter (measured on live specimens), exhibit a reddish-brown coloration, observed in live specimens (Fig. 4A, Appendix Video S1). In general, the pharynx (Figs. 1A–C: ph) is as described by Meixner (1915). It is situated in the anterior third of the body, measuring 115–160 μm (n=2) in length, with a diameter of 92–116 μm (n=2). The mouth opening, prepharyngeal cavity, and pharyngeal lips are ciliated. An external layer of circular muscles surrounds the pharynx bulb, just inside the septum. Approximately 40 radial muscles run between the internal and the external walls. The pharyngeal lumen is covered with a relatively high nucleated epithelium and is surrounded by an inner circular and an outer longitudinal muscle layer. The brain is positioned immediately anterior to the pharynx, and can be recognized as an eosinophilic mass (Figs. 1C and 2A: br).

**Figure 1:**
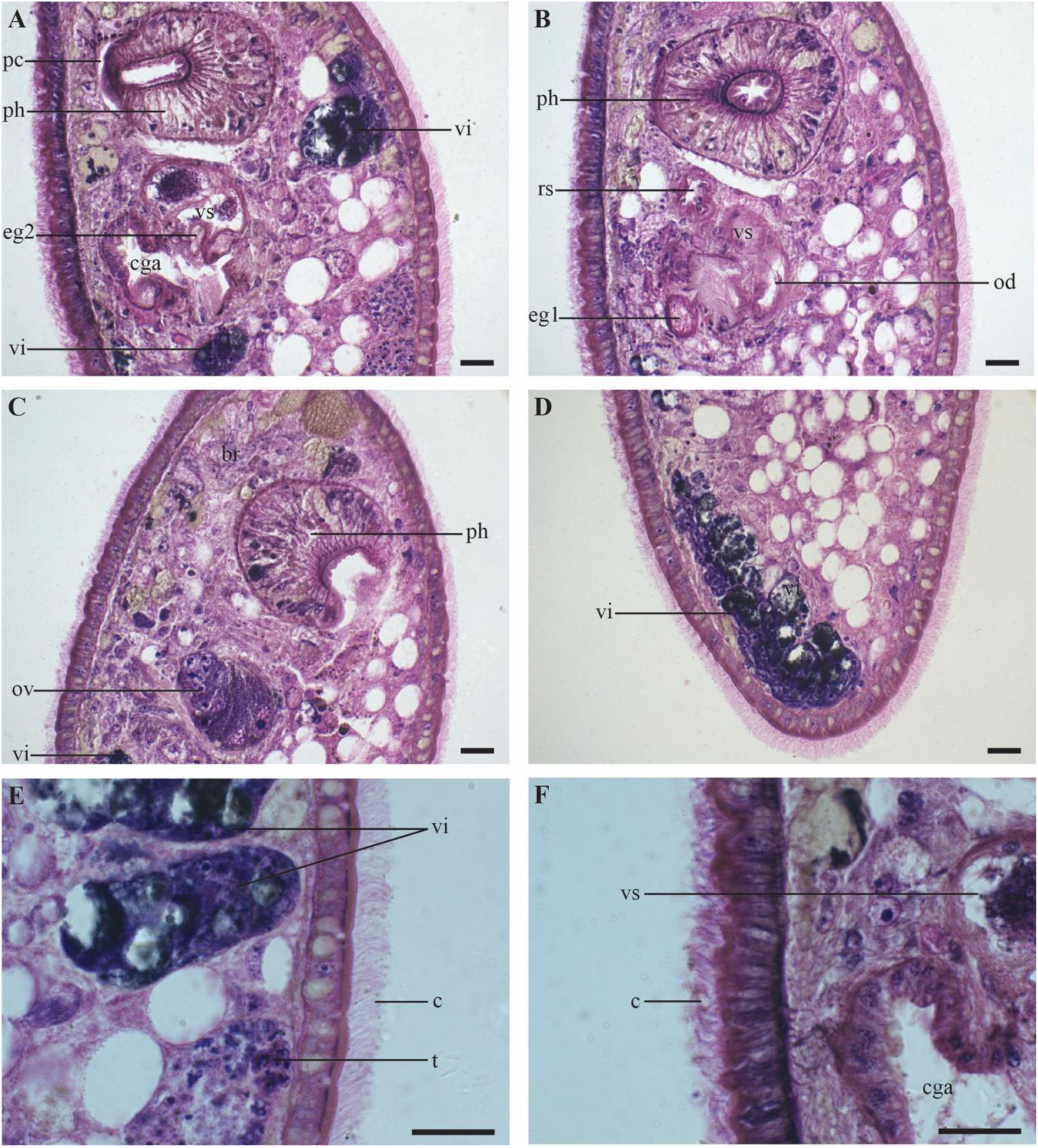
*Strongylostoma simplex*, details of the internal morphology on sagittal sections A-C. Structures oriented with the anterior end toward the top of the plate, D. Posterior end of the body, E. Detail of the dorsal epidermis, F. Detail of the ventral epidermis. Scale bar = 20 µm.

**Figure 2:**
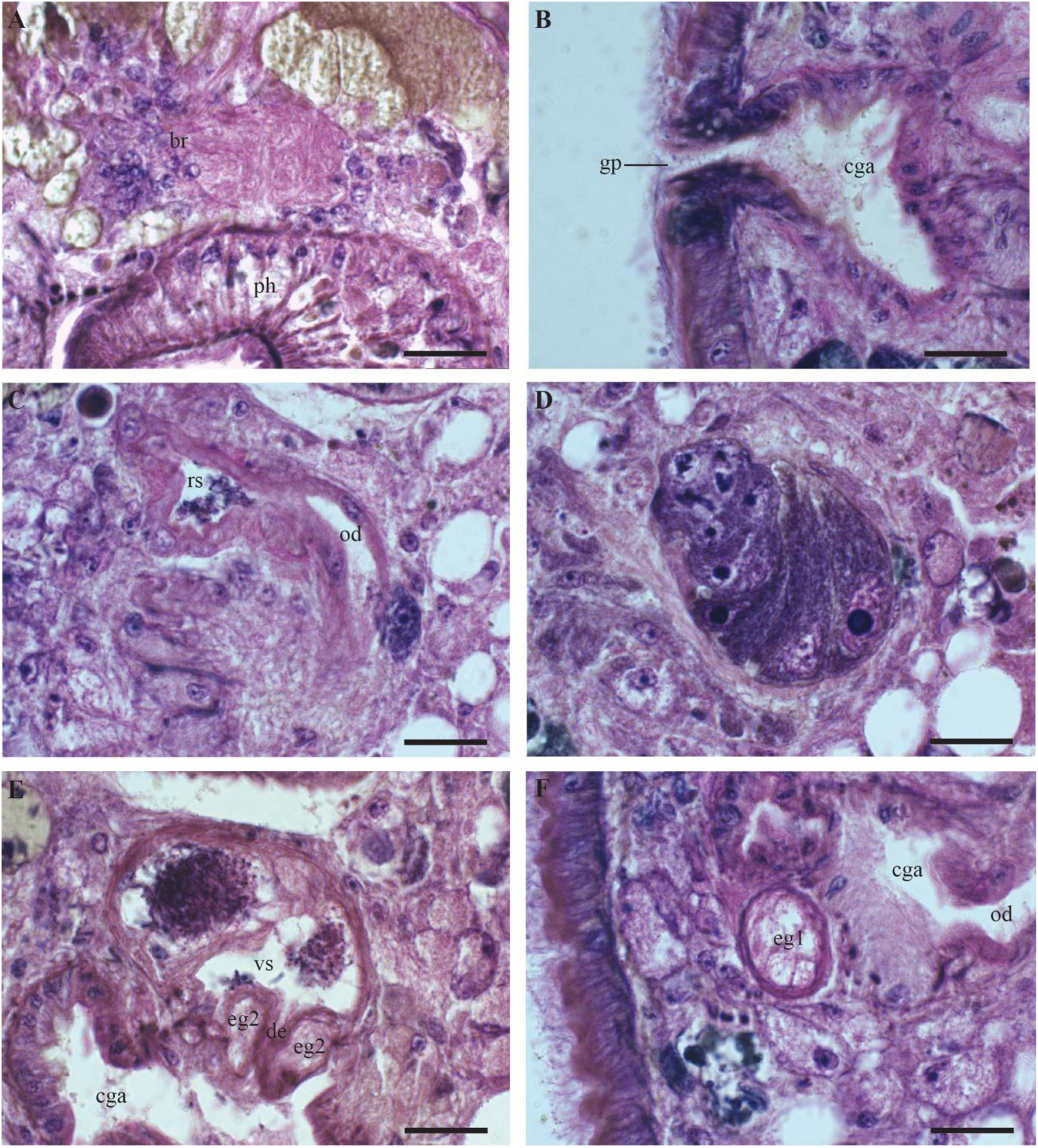
*Strongylostoma simplex*, details of the internal morphology on sagittal sections A. Brain in the anterior part of the body, B. Genital opening and common genital atrium, C. Seminal receptacle with a segment of the oviduct, D. Ovary, E. Seminal vesicle, F. Eosinophilic gland next to the common genital atrium. Scale bar = 20 µm.

Immediately posterior to the pharynx lies the reproductive system, which occupies roughly the middle third of the body (Fig. 3). The ovary (Figs. 1C, 2D, 3A: ov) is inverted pear-shaped and measures 69–71 μm in length (n=2). The oocytes are arranged in a row, with the largest oocytes located most distally. The vitellaria (Figs. 1A, 1D–E, 3A: vi) are dispersed throughout the body, primarily on the dorsal and ventral sides. The vitelloduct (Fig. 3A: vd) is connected to the proximal end of the oviduct, which leads to the seminal receptacle (Figs. 1B, 2C, 3A: rs). The seminal receptacle has a diameter of 15 μm (n=1) and no muscular stalk. A female duct connects the female reproductive system to a common genital atrium (Figs. 1F, 2B, E–F: cga), which measures 52 μm in width and 65 μm in length (n=1). A large eosinophilic gland occurs adjacent to the genital atrium (Figs. 1B, 2F, 3: eg1), though no connection to the genital atrium was found. The genital opening is centrally located and ciliated, with cilia about half the length of those on the epidermis.

**Figure 3:**
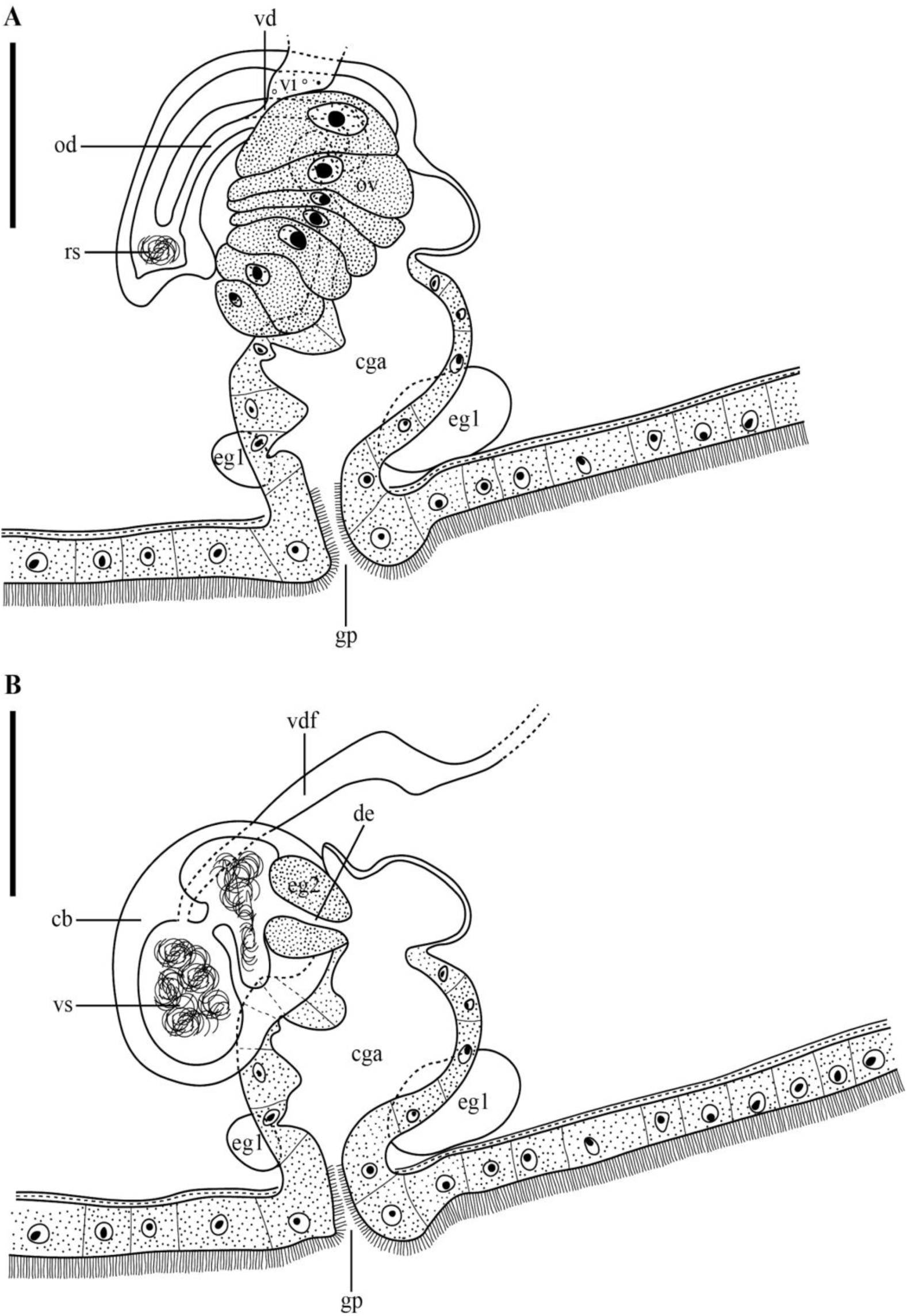
*Strongylostoma simplex* A. Reconstruction of the female reproductive system. The position of the seminal receptacle was slightly moved laterally for visibility purposes. B. Reconstruction of the male reproductive system. Scale bar = 50 µm.

The testes (Figs. 1A and 1E: t) are situated dorsally in the posterior third of the body and vary in shape from oval to rounded, though they are predominantly rounded. The vas deferens (Fig. 3B: vdf) empties into the copulatory bulb (Fig. 3B: cb), which encloses a single seminal vesicle (Figs. 2E, 3B: vs). The seminal vesicle measures 52 μm in width and 68 μm in length (n=1) and is divided into two parts with a minor connection between them. Given the presence of two testes, it is likely that the vas deferens fuse somewhere before entering the bulb. However, we did not observe the separate vas deferens or the point where this fusion may occur. A short ejaculatory duct (Fig. 3A: de) empties into the common genital atrium. It measures 19–25 μm in length (n=2) and is surrounded by a well-developed, eosinophilic prostatic gland (Figs. 1A, 2E, 3B: eg2) with a diameter of 25 μm (n=1), containing a medium-grained secretion. These glands are not mentioned by Meixner (1915), nor by Luther (1963).

#### Remarks

The absence of a proboscis (Willems et al., 2006) and lack of a double connection in the female system (Van Steenkiste & Leander, 2022; Vicente-Hernández et al., 2023) exclude the studied specimens from Kalyptorhynchia and Mariplanellida, respectively, and unambiguously place them within Dalytyphloplanida. The presence of paired, compact testes, a single ovary, follicular vitellaria, a single genital opening, and a pharynx rosulatus positions them within the (paraphyletic) family ‘Typhloplanidae’ (Graff, 1905; Houben, 2013; Houben et al., 2022; Van Steenkiste et al., 2013). This species-rich assemblage comprises 287 described species to date (Tyler et al., 2006–2025), with the specimens under study here designated to *Strongylostoma*.

Species of *Strongylostoma* are characterized by dermal rhabdites, protonephridia that open near the mouth, a genital opening in the anterior two-thirds of the body, a pharynx located in the middle or anterior part of the body, and the absence of a uterus and copulatory atrium (Graff, 1913; Luther, 1904, 1963; Örsted, 1843; Van Steenkiste et al., 2011). These characteristics were corroborated in our studied specimens. Most species of *Strongylostoma* also possess eyes (Örsted, 1843), except for *S*. *coecum* Sekera, 1912. Additionally, most species of *Strongylostoma* typically have a seminal receptacle with a muscular stalk; however, this is not the case for *Strongylostoma simplex simplex* (Meixner, 1915), the (sub)species to which the specimens under study belong.

*Strongylostoma simplex simplex* is morphologically most similar to *S*. *devleeschouweri* Van Steenkiste, Tessens, Krznaric & Artois, 2011. These two species are, for instance, the only ones in the genus lacking spines in the ejaculatory duct and are also distinctive in that the common genital atrium is not divided into two parts—a key feature distinguishing *S*. *simplex simplex* from *S*. *simplex lapponicum* (Luther, 1963; Van Steenkiste et al., 2011). However, *S. devleeschouweri* is distinct from the specimens under study due to its green-spotted coloration, caudally positioned testes, the presence of the genus-typical sphincter around the seminal receptacle stalk, a caudal protrusion of the common genital atrium, and granular eosinophilic prostate glands filling the copulatory bulb (Van Steenkiste et al., 2011).

### Impact of flatworm predation on water fleas

#### Observational findings

During *in situ* field observations, specimens of *Strongylostoma simplex* were observed either in the brood chamber of the water flea *Daphnia magna* (detectable by white reflecting coloration), or free-swimming in the water column (the flatworm is visible to the naked eye). While the observations described below are mainly based on *Daphnia magna*, we also detected flatworm infections in a smaller water flea species that co-occurred in the sampling site, i.e. *Daphnia longispina* (Appendix Fig. S2).

During our detailed observations in the container and Petri dish, we observed flatworms actively chasing the water fleas, and attaching themselves on the carapace (exoskeleton). The swimming speed of the flatworms was visibly faster during chases. Some water fleas were observed to shake the worms off by rapid circular movements (Appendix Video S2). In other cases, flatworms successfully entered the water fleas’ body cavity via the opening in the filter apparatus. Once inside the body cavity, flatworms squeezed themselves into the brood chamber of the water fleas, and moved between the embryos (Fig. 4A, Appendix Video S1). Infected water fleas were occasionally observed to perform ventral flexion of the post-abdomen (Appendix Video S3), a behaviour typical during the release of newborn juveniles (Ebert 2005), here performed possibly as a reaction to the flatworm infection. Additionally, infected water fleas seemed to have reduced swimming performance.

**Figure 4.**
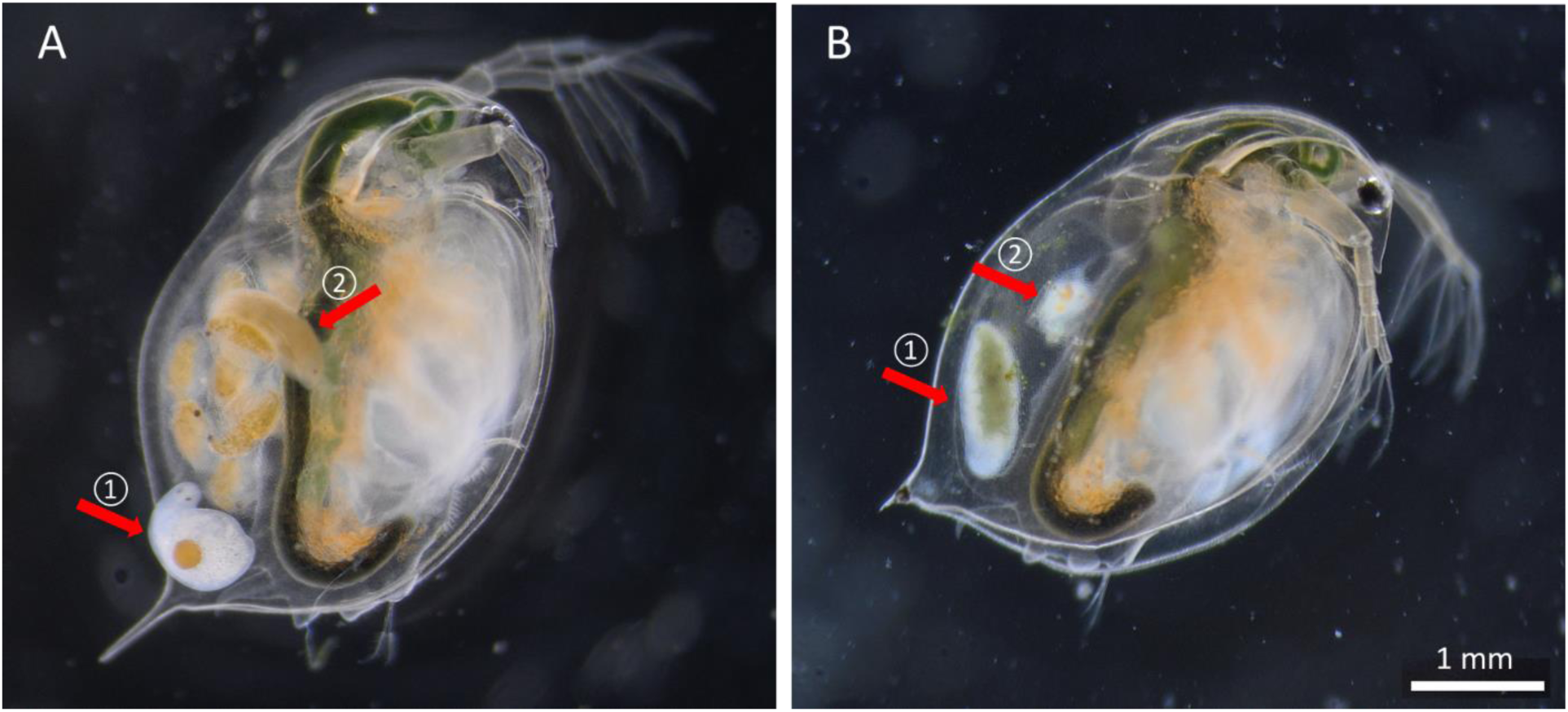
The water flea *Daphnia magna* infected by the flatworm *Strongylostoma simplex*. A) A flatworm, containing an egg, is attached to the water flea carapace (arrow-1), and another flatworm is inside the brood chamber of the water flea, next to the water flea embryos (arrow-2). B) A flatworm is inside an empty brood chamber of the water flea (arrow-1), next to a clump of deformed tissue, possibly belonging to an embryo (arrow-2). Note also the differences in the individual flatworm coloration, possibly due to feeding.

Importantly, we observed partially deformed embryos in flatworm-infected water fleas (Fig. 4B), suggesting a potential brood predation role of the flatworm. While we did not make any direct observation of water flea embryos being eaten by the flatworms, we noted that flatworms found in brood chambers were of darker colouration – in contrast, free-swimming flatworms are more or less white coloured (Fig. 4). This colouration may potentially be due to recently digested water flea eggs. Aside from the potential brood predation behaviour, we also observed flatworms attached to the water fleas’ ovaries and/or midgut, potentially feeding on tissues other than eggs (Appendix Fig. S3).

#### Experimental findings

After 9 days, only 1 out of 8 water fleas survived in the flatworm-treatment, whereas 6 out of the 8 survived in the (flatworm-free) control-group (Fig. 5). This difference was statistically significant (p = 0.0152, Fisher’s exact test).

**Figure 5.**
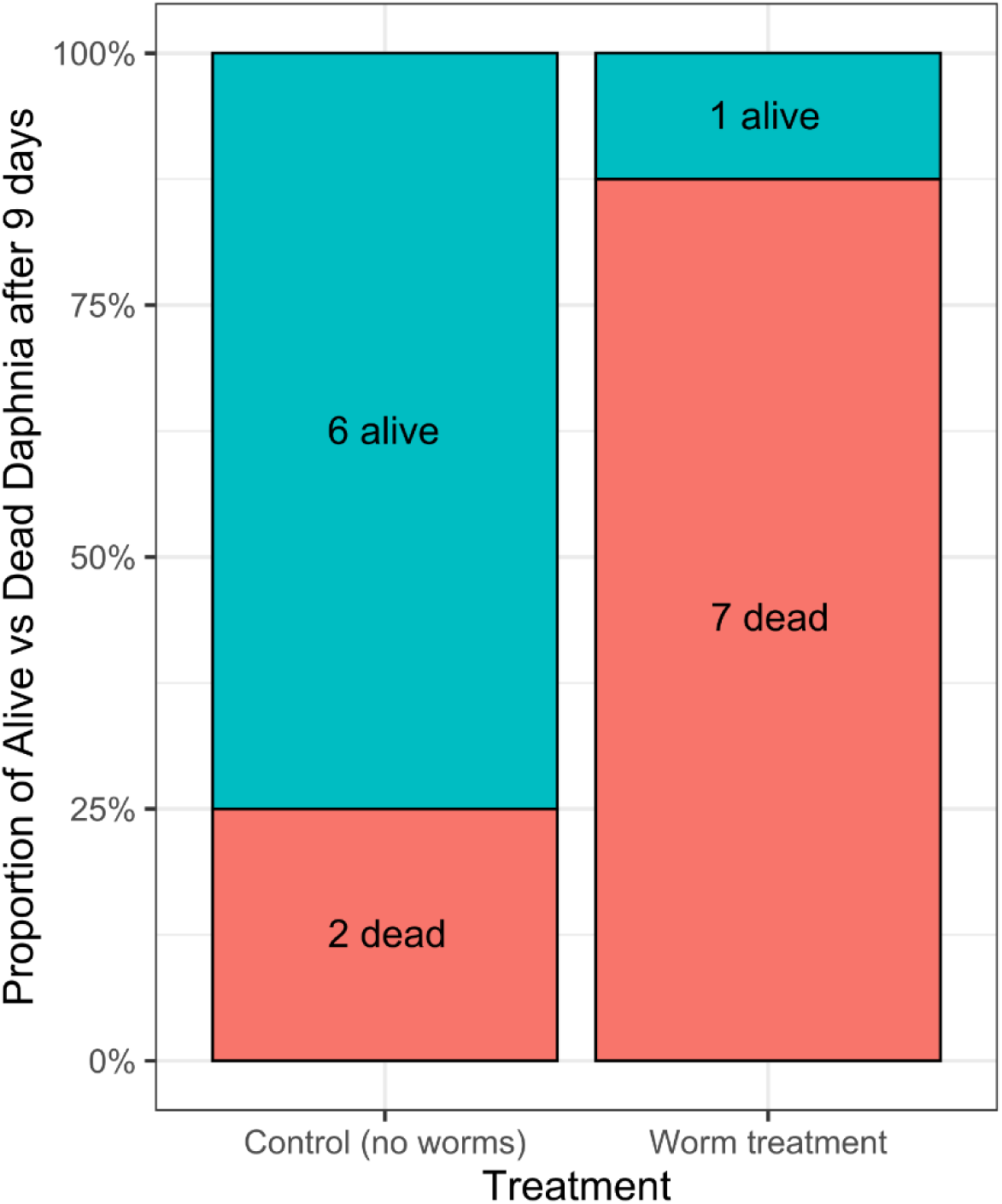
Survival of the water flea *Daphnia magna* after 9 days, either in the absence of flatworms (control group) or presence of flatworms (treatment group).

The average number of offspring produced over 9 days was 14.75 for the control, and 2.25 for the flatworm-treatment group (Fig. 6); a statistically significant difference (p = 0.0424, Wilcoxon rank-sum test). In the flatworm-treatment group, 5 out of the 8 water fleas did not produce any offspring, whereas in the control treatment only 2 water fleas did not produce any offspring (Fig. 6).

**Figure 6.**
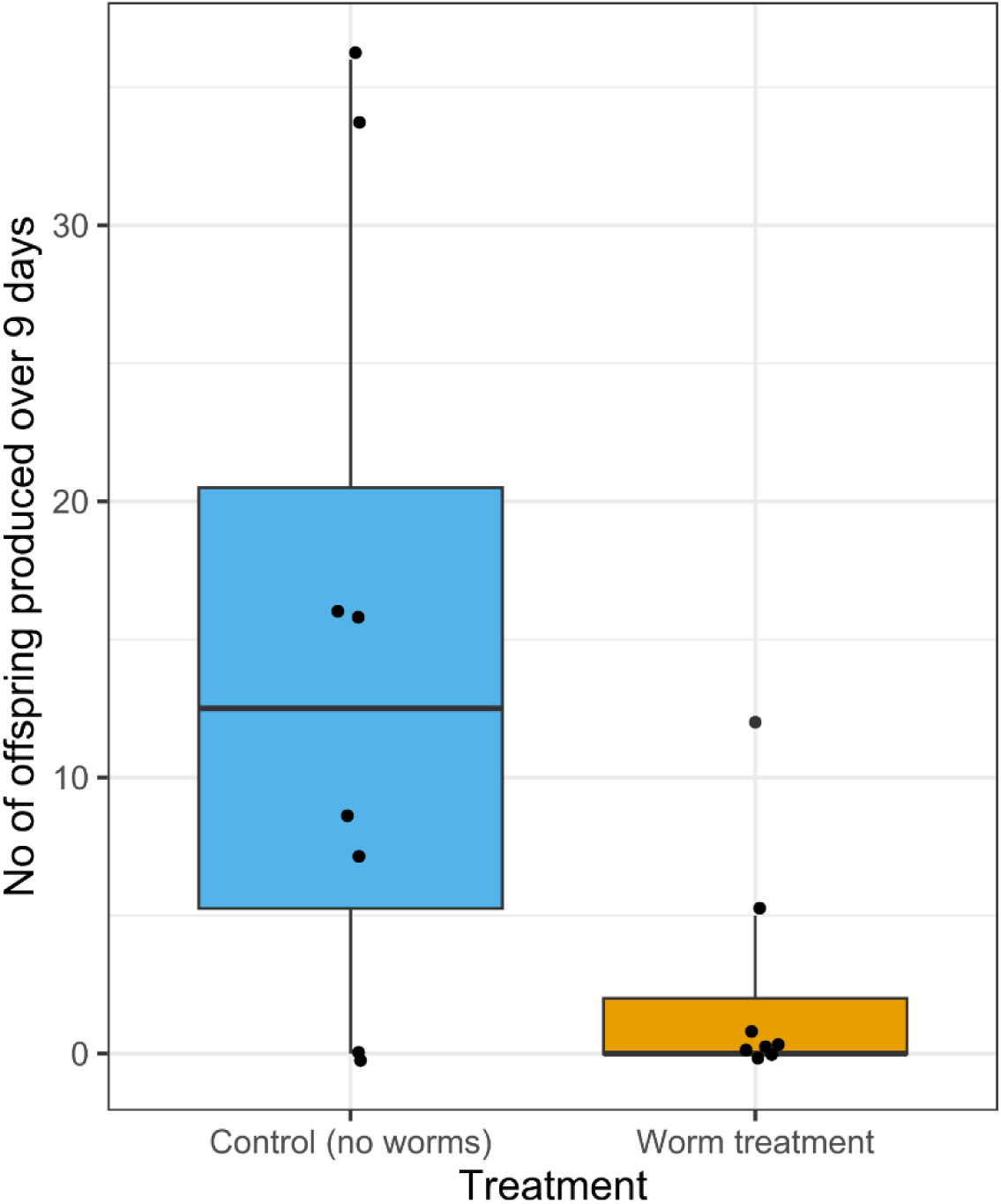
Number of offspring produced by the water flea *Daphnia magna* over 9 days, either in the absence of flatworms (control group) or presence of flatworms (treatment group).

### Conclusions

Our experiment revealed that infection by flatworms caused a drastic reduction in water flea fitness, measured as survival and offspring production. This finding corroborates our visual observations of egg predation by the flatworms, and suggests a strong pressure on water flea populations. Reduced population sizes of zooplankton due to rhabdocoel flatworm predation has been documented (Blaustein & Dumont 1990), including for *Daphnia* (Maly et al., 1981; Wang et al., 2011). Aside from these correlational studies, a laboratory experiment revealed negative effects of the predatory flatworm *Stenostomum leucops* (Catenulida) on the lifespan of the cladoceran *Moina macrocopa* (Nandini & Sarma, 2013). Our finding of reduced survival may be linked to the injury caused by the flatworms when consuming water flea embryos and undeveloped eggs, as well as when transitioning from the body cavity into the brood chamber. Alternative predation methods employed by other flatworms may also be a cause of mortality. The relatively well-studied rhabdocoel flatworm *Mesostoma* employs a wide variety of prey killing mechanisms, including trapping prey in mucus and paralyzing prey via toxins (Blaustein & Dumont, 1990; Dumont et al., 2014). While we did not observe any of these behaviours, at this point we cannot fully exclude them as alternative mechanisms.

Water bodies, including those located in urban areas, provide essential ecosystem services, ranging from protecting biodiversity to recreational and human health benefits (Higgins et al., 2019). Zooplankton grazers such as *Daphnia* are essential organisms in these ecosystems. Algal blooms, which may also come in toxic forms that are dangerous to wildlife, are actively prevented by healthy populations of large grazers such as *Daphnia* (Ger et al., 2016). Reduced population sizes of *Daphnia* in the presence of flatworms may therefore risk the balance of these ecosystems. Importantly, many zooplankton species, including species of *Daphnia,* exhibit diel vertical migration patterns, i.e. grazing on algae close to the water surface at night, and staying close to the bottom to hide from visual predators during day (De Meester et al., 2022). If *Strongylostoma simplex* follows this migration pattern, as was suggested for a species of *Mesostoma* (De Meester & Dumont, 1990), the encounter rate between *Daphnia* and flatworms will be high, resulting in stronger predation pressure on populations of *Daphnia*. Moreover, studies on the predatory flatworm *Mesostoma* reveal higher predation rates on *Daphnia* at warmer temperatures (Beisner et al., 1997; Devkota et al., 2023) and in shallower ponds (Maly et al., 1981). Urban water bodies, which are typically shallow and have higher temperatures compared to rural, natural ponds (Brans et al., 2018), provide habitats that may promote a strong flatworm-predation pressure on *Daphnia*. Light pollution, another anthropogenic stress factor associated with urbanization, can additionally influence this interaction (e.g. for host-parasite interactions in aquatic ecosystems: Poulin, 2023). Finally, our observation that both of the cooccurring species of *Daphnia*, *D. magna* and *D. longispina*, were infected by the flatworm is of concern, as the potential loss of the functional role (i.e. grazing on phytoplankton) of one species may not be compensated by the other.

Based on our discovery of *Strongylostoma simplex* flatworms predating on *Daphnia* water flea embryos, with strong negative effects on the *Daphnia* population, we encourage further research investment into exploring this novel interaction. Later sampling efforts revealed the presence of flatworms in water wells from at least five additional cemeteries in Berlin (data not shown), suggesting this to be a widespread phenomenon in the study region; at least in this type of habitat. So far we have encountered this interaction only in urban cemetery wells, but a potential spread of the flatworms to other water bodies (e.g. urban park ponds, natural lakes) may pose a risk for *Daphnia* populations, hence also the health of aquatic ecosystems. Alternatively, this novel interaction may be restricted to very small water bodies such as cemetery wells (e.g. increased probability of water fleas and flatworms encountering each other due to spatial constraints); a possibility that requires further investigation.

## ACKNOWLEDGMENTS

Dr. Alexander Fürst von Lieven is thanked for valuable discussions on the design and findings of the study. Mrs. Natascha Steffanie is thanked for making the histological sections. NT was supported by the Alexander von Humboldt Research Fellowship and Marie Skłodowska-Curie Actions Fellowship. YLD is supported by the Smithsonian Institution through the Smithsonian Marine Station Postdoctoral Fellowship Program. The research leading to results presented in this publication was carried out with infrastructure funded by EMBRC Belgium - FWO project GOH3817N.

## DATA AVAILABILITY STATEMENT

Data of the experimental trial have been deposited at FigShare: https://doi.org/10.6084/m9.figshare.28380659.v1

## SUPPLEMENTARY MATERIAL

Supplementary figures (https://doi.org/10.6084/m9.figshare.28380935.v1)

Supplementary videos (https://doi.org/10.6084/m9.figshare.28380836.v1)

