## Supplementary Figures for "Tiny killers: first record of rhabdocoel flatworms feeding on water flea embryos"

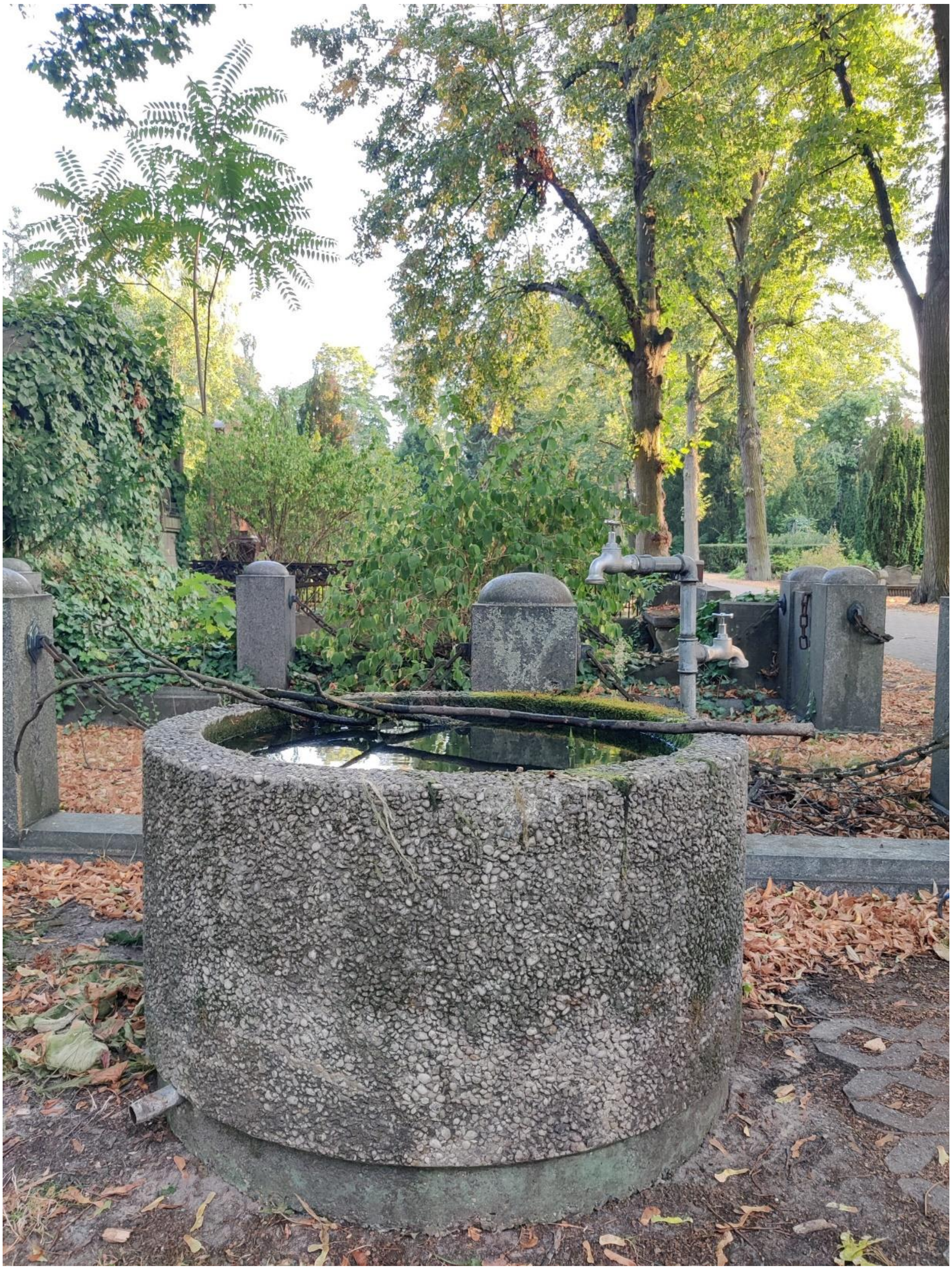

**Figure S1**. The water reservoir in which the observation of *Strongylostoma simplex* flatworms predating on the water flea *Daphnia magna* was first observed (Luisenkirchhof II cemetery, Berlin, Germany).

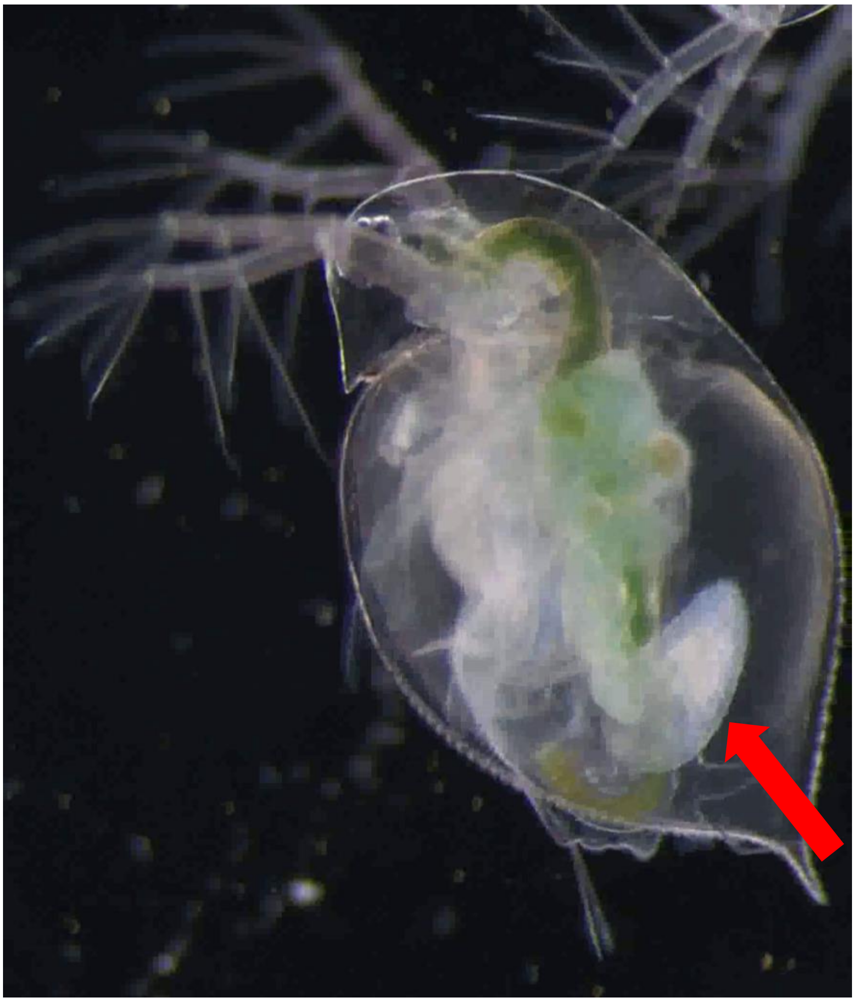

**Figure S2**. Flatworm infection by *Strongylostoma simplex* (indicated with an arrow) in the smaller water flea species, *Daphnia longispina*, that co-occurred in the sampling site with *D. magna*.

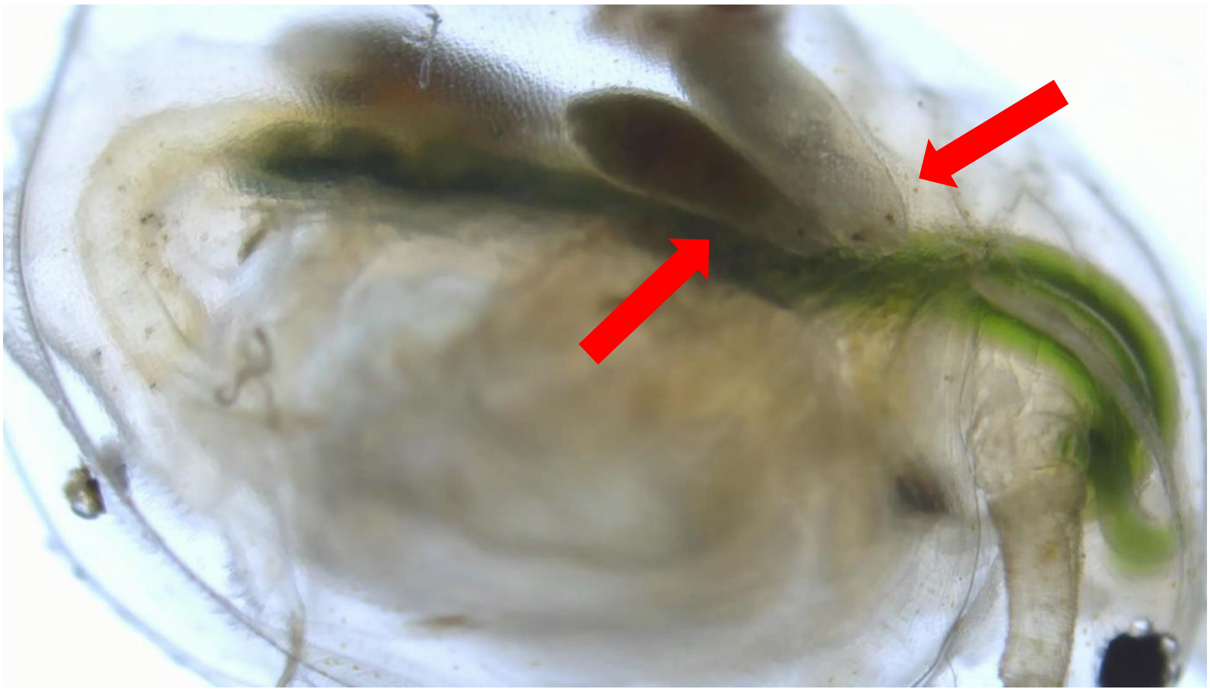

**Figure S3**. Flatworm infection by *Strongylostoma simplex* (indicated with arrows) in the water flea *Daphnia magna*. Note that flatworms seem to be attached to the water fleas' ovaries and/or midgut, potentially indicating feeding behaviour of the flatworms on tissues other than eggs.
